# Small-amplitude head oscillations result from a multimodal head stabilization reflex in hawkmoths

**DOI:** 10.1101/2022.04.24.489321

**Authors:** Payel Chatterjee, Umesh Mohan, Sanjay P. Sane

**Affiliations:** National Centre for Biological Sciences, Tata Institute of Fundamental Research, Bangalore 560065, India

**Keywords:** Negative feedback loop, Multimodal integration, gaze stabilization reflex, head positioning, Oleander hawkmoth *Daphnis nerii*, insect vision, antennal mechanosensation

## Abstract

In flying insects, head stabilization is an important reflex which helps to reduce motion blur during fast aerial maneuvers. This reflex is multimodal and requires the integration of visual and antennal mechanosensory feedback, each operating as a negative-feedback control loop. As in any negative-feedback system, the head stabilization system possesses inherent oscillatory dynamics that depends on the rates and latencies of the sensorimotor components constituting the reflex. Consistent with this expectation, we observed small amplitude oscillations in the head motion (or head wobble) of the Oleander hawkmoth *Daphnis nerii*. We show here that these oscillations emerge from the inherent dynamics of the multimodal reflex that underlies gaze stabilization, and the amplitude of the head wobble is a function of both the visual feedback and antennal mechanosensory feedback from the Johnston’s organs. The head wobble is thus an outcome of a multimodal, dynamically-stabilized head positioning reflex.

## Introduction

Reflexive behaviours in animals rely on feedback from various sensory modalities to achieve control and stability in diverse contexts (1). From a systems perspective, such feedback-control loops possess inherent dynamical properties that depend on the gains and latencies of their sensorimotor components (2). When the gain and latency parameters are affected by disease or injury, the dynamics of feedback-control also changes thereby altering the rhythmic output of these behaviours (3,4). To probe into the constitution of these reflex systems, it is thus useful to adopt a strategy in which we experimentally alter the latencies of the sensory feedback and observe the resulting changes in motor output.

In flying insects, several reflexes constituted as negative feedback loops help to maintain the body posture during flight (1). These reflexes, which are distributed along the body, include the positioning of their antennae (5), head stabilization (6,7), and abdominal flexion (8). Each of these behaviours are necessary features of insect flight, and must be mutually coordinated. In this context, flight-related reflexes in insects are particularly interesting as they are very rapid, and hence push the limits of nervous control. In particular, the head stabilization reflex is crucial for flight due to its intimate association with vision (9). Because insects cannot move their eyes relative to their heads, gaze stabilization is mostly achieved through compensatory head movements which help reduce motion blur of wide-field images during fast aerial maneuvers (7,10).

Head stabilization is also interesting because it involves multimodal sensory feedback. In Diptera, it is mediated by visual feedback from eyes and mechanosensory feedback from halteres (modified hindwings in flies) (7,11) in addition to neck prosternal organs (12). In non-Dipteran insects such as hawkmoths which lack halteres, head stabilization is mediated by visual and antennal mechanosensory feedback (Fig 1A; (9)), and also involve input from prosternal organs although these have not yet been described in moths. The antennal mechanosensory feedback is derived from the Johnston’s organ (JO), which is highly sensitive and spans the pedicel-flagellar joint thus monitoring the relative movements in this joint. This feedback is essential for flight control (13–15). Because visual feedback typically operates at slower timescales than antennal mechanosensory feedback (14), it is not clear how the alteration of individual inputs would affect the head stabilization reflex.

**Figure 1.**
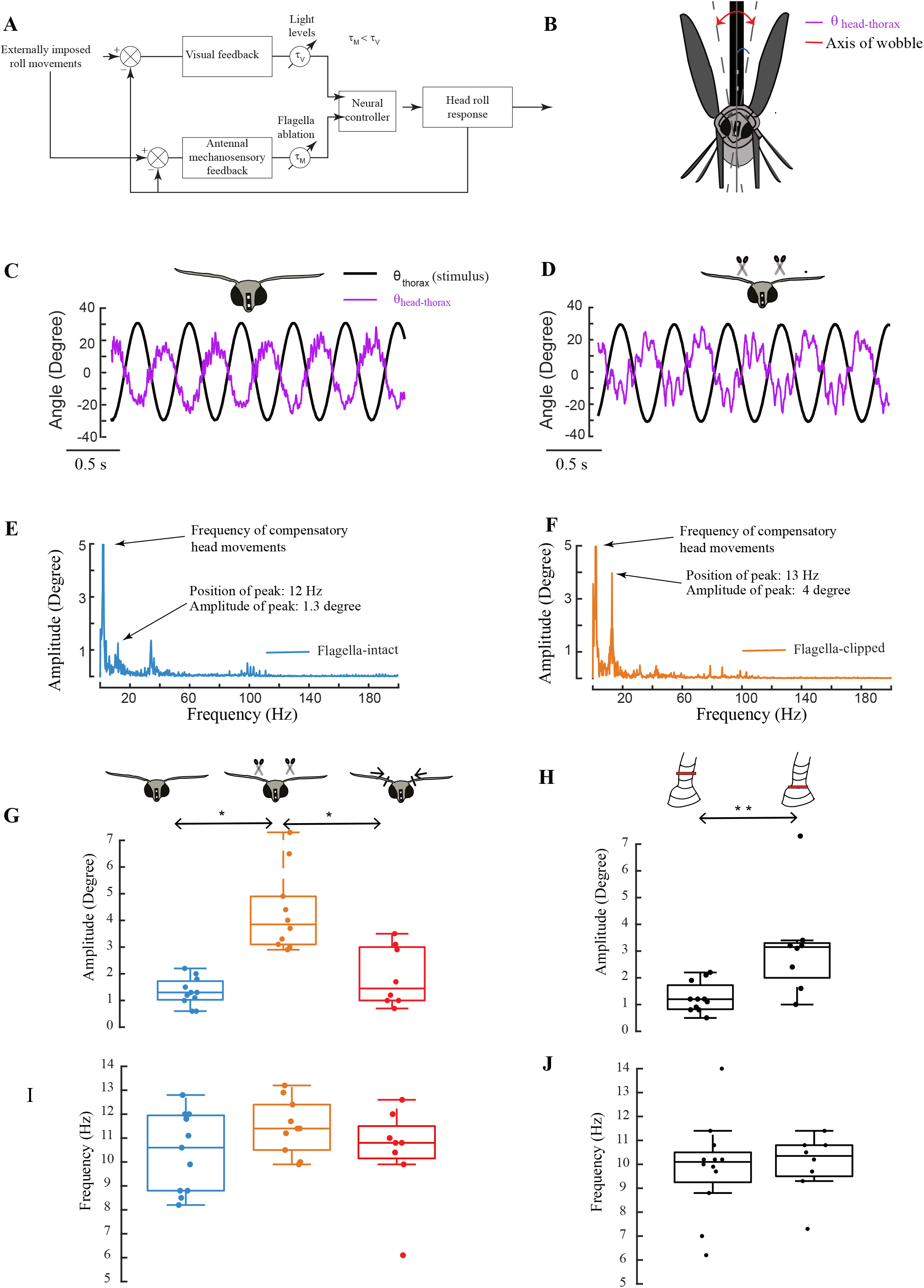
(A) A qualitative model illustrating the feedback loops in head stabilization (9) (B) A schematic illustrating the experimental set-up, head wobble and the angle used for analysis. (C, D) Representative raw plots showing time series of θ_thorax_ (black line) and θ_head-thorax_ (purple line) in presence of imposed roll in the *control* (C) and *flagella-clipped* (D) moths. (E, F) Representative Fourier transforms of the time-series of θ_thorax_ and θ_head-thorax_ in a *control* (blue trace, E) and *flagella-clipped* (orange trace, F) moths. (G-I) Boxplots comparing wobble amplitude (G, H) and frequency (I, J) between *control* (blue box), *flagella-clipped* (orange box*) and flagella-reattached* (red box) moths, and between *sham* (left box, H) and *Johnston’s organ glued* moths (right box; H) in the imposed roll experiments (** depicts p=0.0037 for Wilcoxon Ranksum test). In all box plots, the datapoints are displayed on the boxplots. For comparison in Fig 1G; * signifies p<0.05 using Kruskal-Wallis test followed by Nemenyi test. For comparison in Fig 1H, ** signifies p=0.0037 using Wilcoxon Ranksum test.

As in any feedback-controlled dynamic system, head movements also exhibit oscillations. If these oscillations depend on multisensory feedback, their characteristics should be altered by changes in the feedback from individual sensory modalities. To test these hypotheses, we conducted a series of experiments in tethered Oleander hawkmoths, *Daphnis nerii*. Here, we show that hawkmoths display small-amplitude head oscillations (henceforth called *head wobble*) which depend on both visual and antennal mechanosensory feedback, consistent with our hypothesis. Such oscillatory dynamics emerges naturally from a dynamically stable reflex that determines the head positioning behaviour. When visual feedback is altered by changing light levels or antennal mechanosensory feedback by reducing the flagellar load on the JO, the dynamics of the head movements is altered. Thus, head wobble in hawkmoths provides insights into the dynamics of multimodal reflexes.

## Materials and Methods

### Moth breeding

All experiments were performed on adult Oleander hawkmoths *Daphnis nerii* from our laboratory culture (for husbandry details, see (9)). The choice of *Daphnis nerii* as a study system was motivated by their nocturnal lifestyle, as they can fly stably under very low light levels.

### Treatments

We anesthetized the moths by keeping them in -20°C for ∼8 minutes. The moths were dorsally tethered by attaching a 3mm neodymium magnet with cyanoacrylate glue on the back of their thorax, which was attached to a magnet of opposite polarity on the tether pole. The tether pole was oscillated using a stepper motor to which it was attached (further details in (9)).

### Filming procedure

The moths were filmed from a frontal view with a high-speed camera (V611, Vision research, Ametek, USA) at a frame rate of 1200 frames/second for the imposed roll experiments and at 600 frames/second for the static tether experiments. We used small white markers on black paint on the moth head for contrast and easy digitization. A ring of IR LEDs provided additional illumination during filming.

### Experimental conditions

We recorded the head wobble behavior of moths under two conditions: *imposed roll* and *static tether*.

### Imposed roll stimulus

For the *imposed roll* stimulus, we rotated the tether with a peak-to-peak amplitude of 60 degrees (+/- 30 degrees) at a frequency of 2 Hz (Fig 1B), substantially lower than the 10 Hz typical frequency of head wobble (see results). In the experiments involving *imposed roll* stimulus, the ambient light levels were maintained at ∼250 Lux (twilight). There were 5 experimental groups: *control, flagella-clipped, flagella-reattached, Johnston’s organ (JO) glued* (in which the pedicel-flagellar joint was glued) and its corresponding *sham* procedure in which we applied glue to a few annuli above the pedicel-flagellar joint. These treatments are detailed below.

### Control group

Moths were anesthetized and tethered, but underwent no further manipulations. Thus, their antennal mechanosensory feedback was intact.

### *flagella-clipped* moths

Flagella were excised around the 3rd/4th annulus, drastically reducing the mechanical load on JO and disrupting the antennal mechanosensory feedback.

### *flagella-reattached* moths

Flagella were cut and reattached to their stumps using cyanoacrylate glue. This treatment ensured that only the mechanosensory feedback from JO was restored, whereas feedback from the rest of the antenna remained disrupted due to the severed nerve.

### *JO glued* group/*sham* treatment

Moths were kept on a metal-plate placed atop an ice-bath throughout the treatment procedure (∼30 mins) to render them relatively immobile for the duration of the treatment. In both groups, the head and base of the antennae were descaled to expose the pedicel-flagellar joint, following which cyanoacrylate glue was applied in the pedicel-flagellar joint in the *Johnston’s organ (JO) glued* group, and approximately at the junction between 2^nd^-3^rd^ annulus in the *sham* group to ensure that the gluing procedure itself did not affect head stabilization. By gluing the pedicel-flagellar joint, feedback from JO which spans that joint was severely reduced, and hence similar to moths with clipped-flagella. In contrast, the sham treatment controlled for the effects of the glue application. All the groups were allowed to recover for 30 min post-treatment before starting the experiment.

### Static tether experiments

In the *static tether* case, the tether was kept in a fixed position and the groups were filmed in three light conditions (∼ 200 Lux: clear-sky twilight, ∼ 20 Lux: overcast twilight, < 0.01 Lux: clear night sky/quarter moon). We used a lux meter (Center 337; Range: 0.01-40000 Lux) to measure the ambient light intensity around the moth. The order of the light conditions was randomized across moths. In these experiments, there were two experimental groups: *control* and *flagella-clipped*, prepared as described above.

### Data Analysis

Using custom C++ program in a OpenCV library, we digitized and computed the following angles in MATLAB (also see (9), Fig. 1B)

θ_thorax_ : Angle between thorax and frame vertical

θ_head_ : Angle between head and frame vertical

θ_head-thorax_ : Angle between head and thorax

For imposed roll at 2 Hz, we measured the magnitude of highest peak between 5-15 Hz from the Fourier transforms of θ_head-thorax_ in all the moths. In *static tether* experiments, the stimulus frequency is zero, and hence we computed the magnitude of the highest peak between 2-15 Hz from the Fourier transform.

Details about all statistical comparisons are provided in Supplementary Information.

## Results

### Head wobble amplitude increases when antennal mechanosensory feedback is reduced

In moths that were subjected to an externally imposed roll stimulus, we observed a distinct head wobble in both *control* and *flagella-clipped* moths superimposed upon the compensatory head movements (Fig 1 C-F, G, I). In *control* moths (example in Fig 1C,E), the amplitude of the wobble was 1.3 ± 0.2 degrees (blue box, n = 11; median ± standard error of mean; Fig 1G) and frequency was 10.6 ± 0.5 (blue box, Fig 1I), whereas in *flagella-clipped moths* (example in Fig 1D,F), the wobble amplitude was 3.8 ± 0.5 degrees (orange box, Fig. 1G, n = 10) and frequency was 11.4 ± 0.4 Hz (orange box, Fig. 1I). In moths whose flagella were reattached thereby restoring mechanical load on the JO, the wobble amplitude was 1.4 ± 0.4 degree (red box, n =8, example in Supplementary Fig. 1A, B, Fig. 1G) and frequency was 10.8 ± 0.7 (red box, Fig 1I), similar to the control case. The head wobble amplitude was significantly greater in *flagella-clipped* moths than either *control* or *flagella-reattached* moths (p = 0.0002, Kruskal-Wallis test, followed by Nemenyi test, Fig. 1G). However, the wobble frequencies in the *control, flagella-clipped* and *flagella-reattached* moths were not significantly different from each other (Fig. 1I).

The above experiments showed that antennal mechanosensory feedback is required to minimize head wobble. The key mechanosensory organ in the antenna is the Johnston’s organs, which spans the pedicel-flagellar joint. To determine its role, we glued the pedicel-flagellar joint to reduce JO feedback (*JO-glued* group) and compared these data with a *sham* group in which glue was applied a few annuli above the pedicel-flagellar joint. As in the *flagella-clipped* group, we observed a prominent wobble at 10.4 ± 0.4 Hz in the *JO-glued* group (n = 8, Fig. 1H, J). The wobble amplitude was significantly enhanced in *JO glued* animals (3.2 ± 0.7 degree, n = 8) as compared to the corresponding sham treatment (1.2 ± 0.2 degree, n = 12) (p = 0.0037, Wilcoxon ranksum test; Fig. 1H). The wobble frequencies in both cases were not significantly different (Fig 1J). These data are consistent with the hypothesis that head wobble in hawkmoths depends on mechanosensory feedback from JO (Fig. 1A; (9)).

Such dynamically stable processes naturally generate oscillatory outputs (manifest in this case as a head wobble) due to the gains and delays of the feedback loop associated with each sensory modality. This hypothesis predicts that the reduction of the feedback from the sensory modalities should increase the wobble amplitude, as the system drifts further from its set-point. Consistent with this prediction, wobble amplitude increased when the antennal mechanosensory input was reduced (Fig 1G, H).

### Visual and antennal mechanosensory feedback influence head wobble amplitude

In the experiments described above, we observed the small-amplitude head wobble superimposed with large-amplitude compensatory head movements elicited by an externally imposed roll stimulus which induces optic flow on the retina of the moth. Does head wobble also occur in the absence of the compensatory head movements? Moreover, does head wobble depend on visual feedback in addition to antennal mechanosensory feedback? To address these questions, we conducted a series of experiments on moths attached to a static tether. We altered the latency of visual feedback by conducting the experiments in three ambient light conditions which varied by three orders of magnitude (∼ 200 Lux, ∼20 Lux, < 0.01 Lux) (Fig 2A-F). For each of these light conditions, we repeated the antennal manipulations.

**Figure 2.**
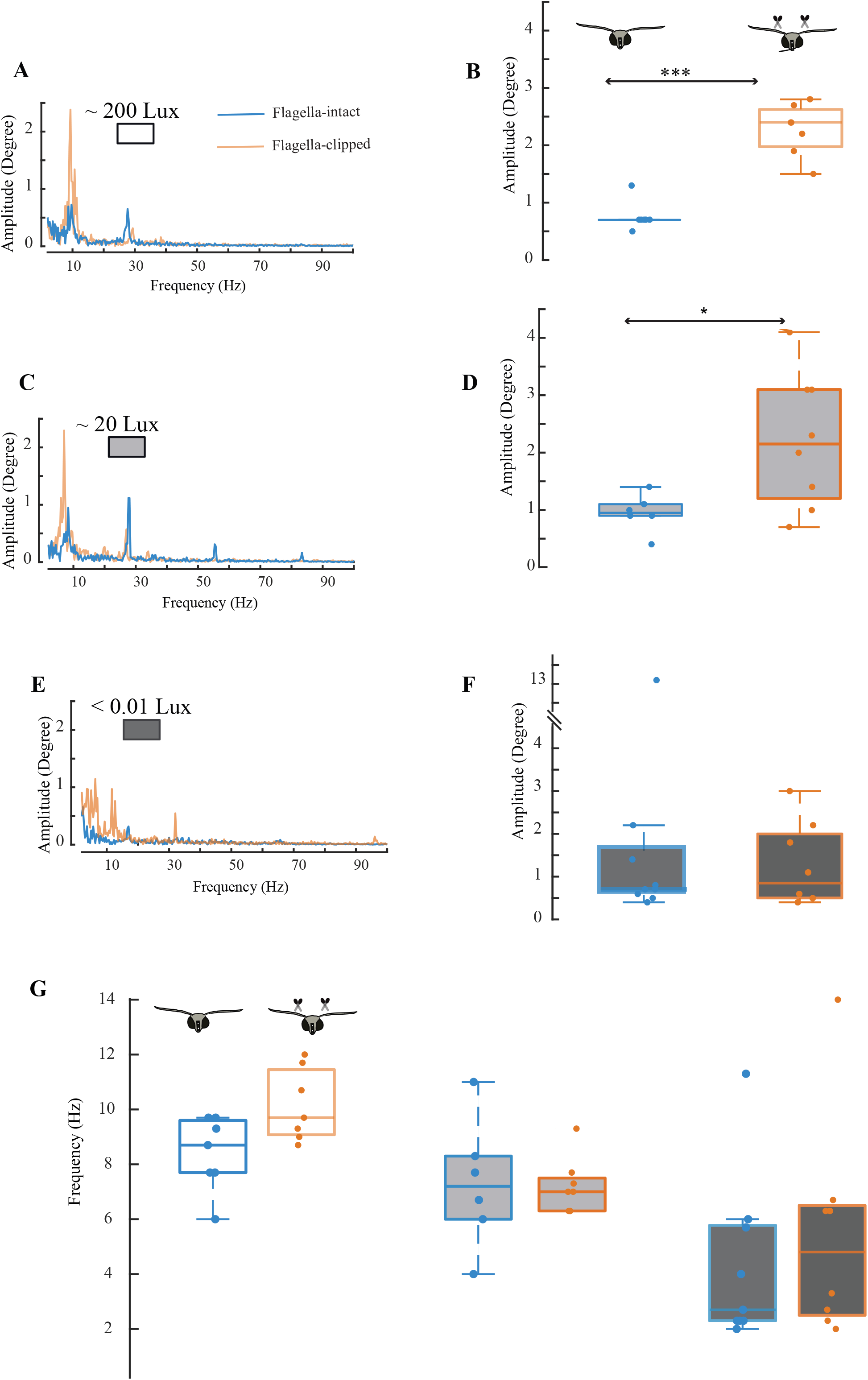
(A-F) Static tether experiments in *control* and *flagella-clipped* moths at ∼ 200 Lux (A-B), ∼20 Lux (C-D), dark (E-F). Representative power spectrums obtained from Fourier transforms of the time-series of θ_thorax_ and θ_head-thorax_ are shown in the left panel for ∼200 Lux (A), 20 Lux (C) and <0.01 Lux (D). In these plots, *Control* moths are shown in the blue trace and *flagella-clipped* moths in orange trace. Boxplots in the right panel compare wobble amplitude between *control* (blue box) and *flagella-clipped* moths (orange box) at ∼ 200 Lux (B, white), ∼ 20 Lux (D, grey), dark (F, black). (G) Box plot comparing wobble frequency between *control* and *flagella-clipped* conditions at the three light levels. For Fig 2B, *** signifies p=0.0006 by Wilcoxon Ranksum test. For Fig 2D * signifies p=0.393 by Wilcoxon Ranksum test.

At an ambient light intensity of ∼200 Lux, the wobble amplitude in *control* moths (blue box, 0.7 ± 0.1, n=7) was lower than in *flagella-clipped* moths (orange box, 2.4 ± 0.2 deg, n=7; p= 0.0006, Wilcoxon ranksum test) (Fig 2 A, B). Thus, antennal mechanosensory feedback helps minimize head wobble amplitude at ∼200 Lux. We observed a similar result at light intensity of ∼20 Lux (Fig 2 C, D) in which the wobble amplitude in *flagella-clipped* moths (orange box, 2.2 ± 0.4 deg, n=8) exceeded the value in *control* moths (blue box, 1 ± 0.1 deg, n=6; p=0.0393, Wilcoxon ranksum test). At 20 Lux, the variability of head wobble amplitude values was greater for *flagella-clipped* moths, suggesting that they had a greater difficulty in minimizing head wobble, perhaps due to the increased latency in visual feedback combined with reduction of antennal mechanosensory feedback.

Under dark conditions (< 0.01 Lux), the wobble amplitude in *flagella-clipped* moths (0.8 ± 0.3 deg, n=8) was similar to *control* moths (0.7 ± 1.4 deg, n=9) (p=0.9125, Wilcoxon ranksum test) (Fig. 2E, F). In the *flagella-clipped* moths, wobble amplitude in dark conditions was significantly lower than at ∼20 Lux (p<0.05, Friedman test, followed by Bonferroni correction, Supplementary Fig. 1C, Fig. 2C-F), but the wobble amplitudes at ∼ 200 Lux and ∼ 20 Lux were similar (∼ 200 Lux: 2.4 ± 0.2 deg, n=7; ∼20 Lux: 2.2 ± 0.4 deg, n=7).

Is wobble frequency a function of these multimodal inputs? At all the light levels, the frequency of head wobble was similar between *control* (∼200 Lux: 8.7 ± 0.5 Hz, ∼20 Lux: 7.2 ± 1 Hz, dark: 2.7 ± 1) and *flagella-clipped* (∼200 Lux: 9.7 ± 0.5 Hz, ∼20 Lux: 7 ± 0.4 Hz, dark: 4.8 ± 1.4) moths (Fig. 2G). We hypothesized that wobble frequency will decrease as light intensity decreases, due to increased latency of the visual feedback. To test this, we compared the *flagella-clipped* moths for which trials of all light levels were present, because they likely depend on visual feedback. Consistent with the hypothesis, wobble frequency slightly reduced when light levels went from ∼200 Lux (9.7 Hz ± 0.4 Hz, n=7) to ∼ 20 Lux (7 ± 0.4 Hz, n=7) but the reduction was greater when the illumination was <0.01 Lux (6.3 ± 1.5, Supplementary Fig. 1D, p < 0.05, Friedman test followed by Bonferroni correction). Thus, the experiments on static tether also highlight the importance of both visual and antennal mechanosensory feedback for minimizing head wobble.

## Discussion

Oscillatory dynamics is a natural consequence of negative-feedback processes in diverse biological systems, including those involved in stabilizing motor output using sensory feedback (1). Such oscillations have been observed and studied in many different systems ranging from postural sway in humans (16), postural ankle oscillations during constant heel elevation (17), slow eye oscillations in humans (18), dynamic whole-body oscillations in weakly electric fish (19,20), maintaining constant hand and finger position in humans (21,22), and hover-feeding from flowers in hawkmoths (15). In all these cases, the dynamical output changes when the latencies of the sensory or motor components of the system undergo an alteration, induced either experimentally (e.g. 15, 23) or due to disease or injury (e.g. 24).

Our study shows that head wobble in moths is an outcome of the dynamics of a multisensory head stabilization reflex (Fig. 1A). In Diptera, mechanosensory feedback for head stabilization is derived from halteres as well as the prosternal organ. Although head oscillations are reported in Diptera, the role of haltere and visual inputs is unclear. For instance, in tethered blowflies, head fluctuations of ±10 degrees occur in absence of an imposed roll stimulus in a homogenous visual background (6). Small-amplitude head oscillations are also observed in *Drosophila* at a frequency of ∼15 Hz in presence of static visual features (25). In hawkmoths, halteres are absent and the prosternal organ is not yet described. However, previous studies show antennal mechanosensory feedback from the JO is crucial for flight (13-15) as well as head stabilization (9). This study renders further support to the role of antennal mechanosensory feedback in maintaining head position. When antennal mechanosensory feedback from JO was altered, hawkmoths faced difficulties in maintaining their aerial position, as manifest from whole-body oscillations in a flower tracking assay in the diurnal hawkmoth, *Macroglossum stellatarium* (15). The amplitude of jitter during hover-feeding in *flagella-clipped* moths was greater than in *flagella-intact* and *flagella-reattached* conditions, similar to the wobble amplitude in *Daphnis nerii*. In these experiments, the wobble or jitter emerges as the nervous system tries to maintain a stable state using sensory feedback. Altering the latency of these inputs changes the head wobble characteristics in a manner that is consistent with the hypothesis presented here.

## Acknowledgements

M. Kemparaju and Allan Francis Joy helped maintain the moth culture, Shivansh Dave and the mechanical and electronics workshops helped with the experimental set up. Funding for this study was provided by grants from Air Force Office of Scientific Research (AFOSR) # FA2386-11-1-4057 & # FA9550-16-1-0155, & National Centre for Biological Sciences (Tata Institute of Fundamental Research) to SPS. We acknowledge support of the Department of Atomic Energy, Government of India, under project no. 12-R&D-TFR-5.04-0800 and NCBS core computational facilities, supported under 12-R&D-TFR-5.04-0900.

## Author contributions

Study design: S.P.S and P.C Methodology: U.M, P.C, Experiments and analysis: P.C Manuscript preparation: P.C, U.M., S.P.S

## FIGURE LEGENDS

**Supplementary Figure 1**. (A) Representative raw plots showing time series of θ_thorax_ (black trace) and θ_head-thorax_ (purple trace) in presence of imposed roll in *flagella-reattached* moths. (B) Representative power spectrums obtained from Fourier transforms of the time-series of θ_thorax_ and θ_head-thora_ in *flagella-reattached* moths (red trace). (C, D) Boxplots comparing wobble amplitude (C) and frequency (D) in *flagella-clipped* moths between ∼200 Lux, ∼20 Lux and <0.01 Lux. For a comparison of the moths between light levels, we used a subset of moths from Fig 2 which underwent trials for all light levels. For these moths, we could statistically compare between the frequencies and amplitudes of their head wobble as a function of light levels. For Supp Fig 1 A and B, * signifies p<0.05 Friedman’s test followed by Bonferroni correction.

