## Supplementary File and Figure for "Small-amplitude head oscillations result from a multimodal head stabilization reflex in hawkmoths"

### Supplementary Information

#### Details about statistical comparisons

1. For comparison of independent samples between two groups, we used a non-parametric test: Wilcoxon rank-sum. The treatments: Sham, JO-glued, Control, Flagella-clipped were conducted on different groups and therefore, we considered them as independent samples (Fig. 1 H, J,B,D,F,G).
2. For comparison of independent samples between three groups, we used a non-parametric test, Kruskal-Wallis. We used a post-hoc Nemenyi test for pair-wise comparison. Because the treatments (*Control, Flagella-clipped, Flagella-reattached*) were conducted on different moths, we considered them as independent samples (Fig. 1 G,I).
3. For comparison of dependent samples between three groups, we used a non-parametric test, Friedman's test. We used a post-hoc Bonferroni correction for pair-wise comparison. Because the same group went through the different light levels, we considered them as dependent samples (Supplementary Fig. 2C,D).

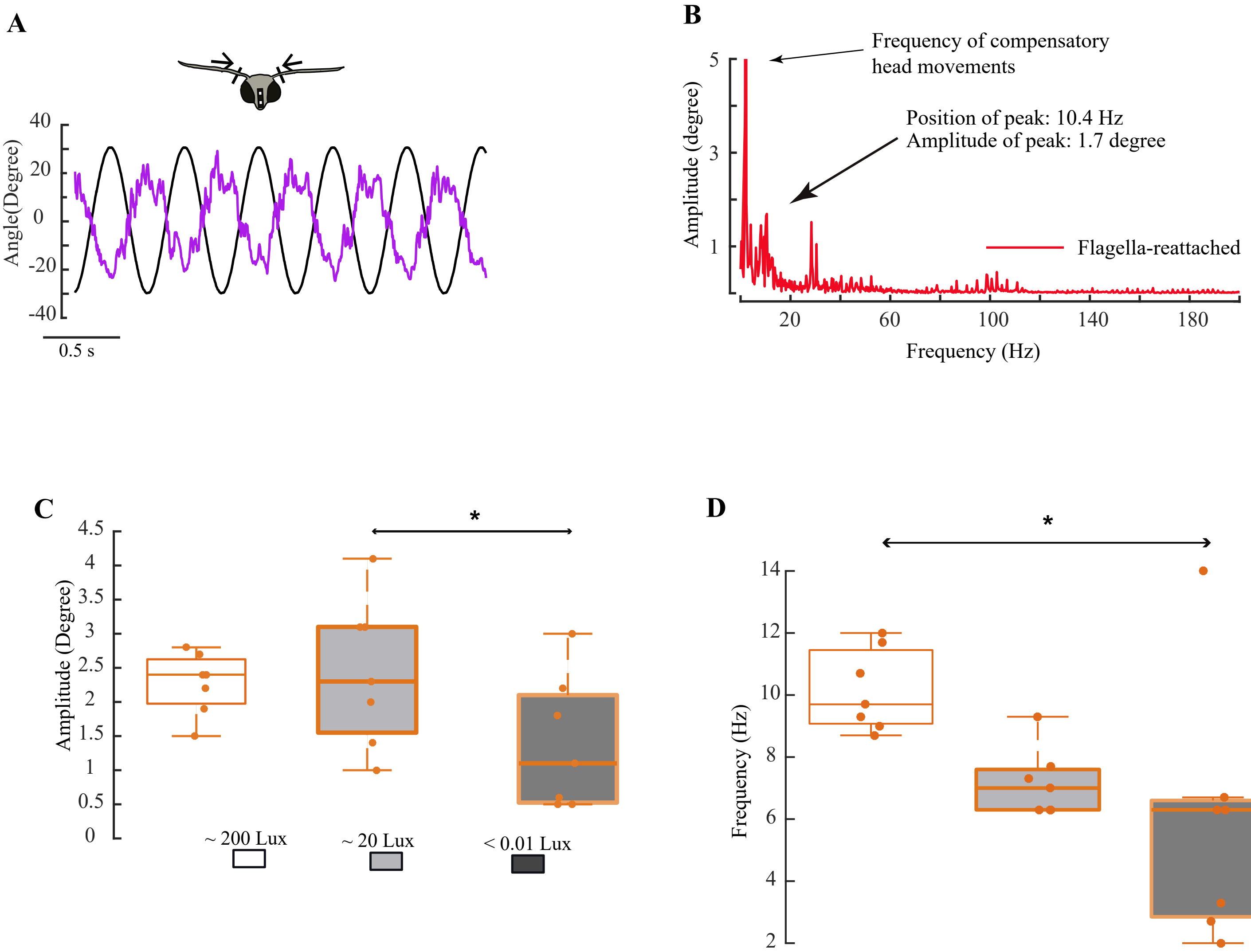
A
Angle(Degree)
40
20
0
-20
-40
0.5 s
B
Frequency of compensatory head movements
Position of peak: 10.4 Hz
Amplitude of peak: 1.7 degree
Amplitude (degree)
Flagella-reattached
Frequency (Hz)
C
*
Amplitude (Degree)
~ 200 Lux
~ 20 Lux
< 0.01 Lux
D
*
Frequency (Hz)
